# Exploiting protein language model sequence representations for repeat detection

**DOI:** 10.1101/2024.06.07.596093

**Authors:** Kaiyu Qiu, Stanislaw Dunin-Horkawicz, Andrei Lupas

## Abstract

Duplication is an essential evolutionary mechanism that operates at the scale of chromosomes, large chunks of DNA sequences, genes, protein domains, and shorter motifs. The study of duplication is central to understanding protein evolution, but the detection of repetitive sequence patterns is often challenging due to decreasing similarity between internal repeats resulting from long-term divergence. The most sensitive sequence-based repeat detection method, HHrepID, relies on the construction of multiple sequence alignments (MSAs) to enhance homology signals and thus facilitate the detection of very ancient duplications. However, such an alignment-based approach is slow and limits the ability to perform large-scale scans. Recent advances in protein representation learning have introduced sequence embeddings extracted from protein language models as a powerful and much faster alternative to MSAs. Protein sequence representations have been shown to be effective in homology detection, as exemplified by software such as our recently developed pLM-BLAST. In this study, we implement pLM-Repeat, a pipeline built upon pLM-BLAST, to identify repeats encoded in sequence embeddings. pLM-Repeat achieves comparable sensitivity to HHrepID in detecting the presence of repeats, while predicting many more repeat units and providing significantly better run times. We also trained an auxiliary neural network, DeepRepeat, to detect domains with patterns similar to well-characterized repeat folds to support rapid filtering. Using our newly developed tools, we scanned the AFDB90v4 database and identified a collection of novel and undescribed repeat domains.

## Introduction

The first discovery of duplicated sequence patterns in proteins dates back to 1966 when Dayhoff and colleagues identified a repeat pattern in a 55-residue ferredoxin (1). They speculated that the protein might be an ancient relic given its repetitive nature and fundamental metabolic function. Over time, proteins with repetitive amino acid patterns have been found to be prevalent in all domains of life. Genomic analysis more than two decades ago revealed that tandem repeats occur in 14% of known proteins and are particularly enriched in eukaryotes (2). A recent survey revisiting the UniProtKB database estimated that more than 50% of deposited proteins contain at least two repeats (3). Proteins with repetitive sequences adopt structures that are typically stabilized by interactions between individual repeat units and exhibit a wide range of structural diversity. This diversity forms the basis for classification hierarchies of different repetitive folds. For example, Kajava proposed a classification scheme of repeat proteins based on both topological features and length of repeat units (4). This framework captures a wide range of repeats, from crystalline aggregates with 1-2 residues per repeat to fibrous repeats such as collagens and coiled coils stabilized by interchain interactions. It also includes elongated repeat folds consisting of tandem arrays packed together to form long and open structures, as well as closed repeat folds where the first and last units are close enough to form a circular topology. Finally, it includes beads-on-a-string proteins formed with repeat domains that can fold independently. This classification system has been adopted by many repeat protein databases, such as RepeatsDB (5), the most comprehensive repository of repeat protein structures curated from the PDB database.

Because of their inherent repetitiveness, repeat proteins provide a versatile platform for a wide variety of studies. For example, their symmetric nature makes them an ideal model system for studying cooperativity in protein folding (6). In addition, protein tandem repeats are involved in various biological processes, such as maintaining structural integrity, mediating interactions, and regulating cascade pathways, and are therefore often implicated in disease (7). From an evolutionary perspective, repeat proteins play a central role in investigating how modern proteins evolved from ancestral peptides, as duplication is recognized as the primary driver of molecular novelty (8). Finally, the success of natural repeat proteins inspires scientists to design artificial repeat proteins in a programmable manner to implement specific functionalities (9).

Given the importance of repeat proteins, how to detect them remains a classic and crucial bioinformatics task, typically approached via sequence-based and structure-based methods (10). Sequence-based algorithms can be further classified into several types. Methods based on short strings are adept at detecting short and highly repetitive tandem repeats, including XSTREAM using string extension (11) and T-REKS using k-means clustering (12). Fourier transform analysis is another algorithm for detecting periodicity within sequences, but struggles with short repeats (13). Alternatively, methods such as TPRpred use prior knowledge by searching query sequences against pre-computed profiles for repeat detection (14). However, the most efficient algorithm for identifying *de novo* repeats, including imperfect and long repeats, is self-sequence alignment (SSA), i.e., aligning the protein sequence of interest to itself using the Smith-Waterman (SW) algorithm and inferring internal repeats from the resulting sub-optimal local alignments. SSA-based software such as RADAR typically receives a single sequence as input (15). Similarly, HHrepID takes a single sequence as input, but then uses it to find homologs and build a multiple sequence alignment, from which suboptimal alignments are inferred by profile-profile self-comparison (16). The use of MSAs makes HHrepID to stand out as the most sensitive sequence-based method over the years, but it reduces its speed and applicability to large scans.

In addition to sequence-based approaches, tandem repeats can also be detected at the structural level by various algorithms, such as Console, which is based on contact map analysis and template matching (17), CE-symm, which uses structural self-alignment and order detection (18), and SymD, which detects internal symmetry by aligning circular permuted structures to their original forms (19). Structure-based approaches generally outperform sequence-based ones because sequences evolve faster than structures, so sequence repeats are less conserved and more difficult to detect. Despite their effectiveness, however, structure-based methods are limited by the availability of 3D structures. Although advances in structure prediction are improving accessibility, sequence data remain more abundant, underscoring the need for continued development of sequence-based methods. It is also important to note that significant sequence similarity typically reflects common ancestry, i.e., homology, whereas structural similarity, even significant, may be the result of convergent evolution. Therefore, sequence-based repeat detection is more informative for studies aimed at understanding protein evolution.

In the search for new ways to detect protein repeats, we focused on the possibilities offered by protein language models (pLMs). These models, inspired by natural language processing techniques, are specialized deep learning models that are trained in a self-supervised manner (e.g., masked language modeling) to understand the grammar of protein sequences. In recent years, numerical representations extracted from pLMs, such as SeqVec (20), ProtT5-XL-U50-half (ProtT5) (21), and Evolutionary Scale Model (ESM) (22), have illuminated various downstream applications for protein science, including antibody design (23), protein-protein interaction prediction (24), and other transfer learning tasks such as detection of transmembrane regions (25) and signal peptides (26). Interestingly, protein representations also contribute to applications involving sequence searching and processing, such as building MSAs (27), predicting structural similarity from sequences (28), and converting sequences into profiles/3Di sequences for further searching (29). We recently developed pLM-BLAST, a homology detection tool based on direct comparison of sequence embeddings (30). pLM-BLAST replaces the fixed substitution matrix (e.g., BLOSUM62) with a context-dependent similarity matrix composed of the similarity of each individual residue pair. This context-dependent substitution matrix is used to construct a score matrix, which is then subjected to the traceback procedure adapted from the Smith-Waterman (SW) algorithm. Notably, pLM-BLAST traces back from all positions in the score matrix, not just from the cell with the highest score as in the original SW algorithm, to effectively report all significant traces. While pLM-BLAST offers good speed due to its independence from the extensive sequence database searches required for MSA generation, it shows performance comparable to HHpred (31), a highly sensitive method based on MSAs, and even detects remote homology signals that HHpred may not capture.

The success of pLM-BLAST and similar homology detection methods makes us wonder if protein representations could also be used for repeat identification. In this study, we used sequence embeddings extracted from protein language models to develop a repeat detection procedure, called pLM-Repeat, based on our recent homology detection software pLM-BLAST. In our benchmarks, pLM-Repeat shows promising performance compared to the most sensitive sequence-based method, HHrepID, while providing significantly faster analysis. To improve the applicability of pLM-Repeat in large, high-throughput scans, we trained an auxiliary neural network to rapidly detect potentially repetitive regions with patterns similar to those seen in repeat protein domains deposited in current databases. As an application of our tools in the era of ever-expanding databases, we scanned 682,563 sequences from the AlphaFold Database (AFDB) (32) and identified 5460 repeat domains displaying novelty.

## Methods

### Dataset

The RepeatsDB database (version 3.2, https://repeatsdb.bio.unipd.it/) was used to generate a set of repeat proteins filtered to 90% sequence identity and 80% coverage using MMseqs2 (version 13.4511) (33), resulting in a dataset of 2056 repeat proteins. Both sequences and structures of the PDB entries of this dataset were retrieved using scripts provided on the PDB website (https://www.rcsb.org/downloads/). To construct a dataset of proteins without repeats, we adopted the protocol described by Alvarez-Carreño et al (34). First, we clustered PDB chain sequences at identity and coverage cutoffs of 30% and 80%, respectively, using MMseqs2. Then, entries marked as repetitive were excluded based on annotations from the InterPro database (35). Finally, the remaining structures were evaluated for the presence of internal symmetry using the SymD software (version 1.61) (19). Proteins with a Z-score of 4 or less were considered to be non-symmetric and thus retained, resulting in a dataset of 8710 non-repetitive proteins. From this set, we randomly selected 2100 proteins to create a negative dataset of comparable size to the positive dataset. These proteins were analyzed using HHrepID at a range of P-values, and those reported as repetitive were manually reviewed and removed if they showed repeat patterns at the sequence or structure level, resulting in a total of 1987 non-repetitive proteins. Both the repetitive and non-repetitive protein datasets served as benchmark sets for pLM-Repeat and as neural network training data (see sections below).

### Implementation of pLM-Repeat

The pLM-Repeat procedure includes the steps described below, with an example of domain 2QJ6_A showing intermediate outputs in Fig. 1.

**Figure 1.**
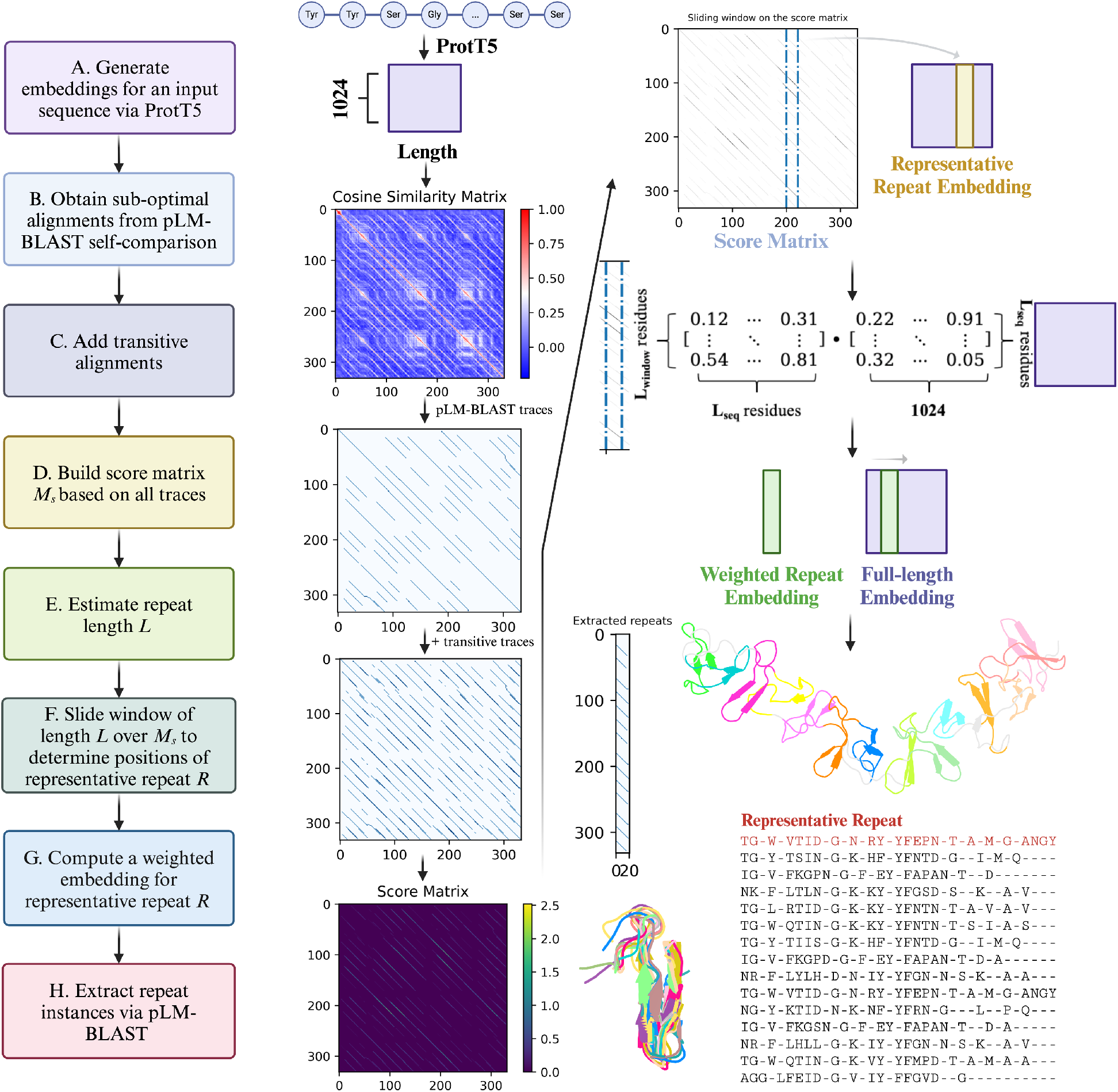
pLM-Repeat pipeline. The main steps of pLM-Repeat are shown as a flowchart, and the main intermediate outputs of the pLM-Repeat workflow are presented using the domain 2QJ6_A as an example. The workflow of pLM-Repeat includes (A) generating protein embeddings with the ProtT5 model, (B) retrieving suboptimal alignments by pLM-BLAST self-comparison, (C) enriching the alignment matrix by applying transitivity, (D) constructing the score matrix from collected traces, (E) estimating the repeat length, (F) determining the positions of the representative repeat, (G) computing a weighted repeat embedding, and (H) extracting repeat instances.

1. The input sequence of length *Lseq* is first embedded as a residue-wise vector of shape (*Lseq*, 1024) by the ProtT5 model (21). This embedding is then passed to pLM-BLAST for self-comparison in local mode with the following parameters: a window length of 15, a minimum span length of 15, a Sigma factor of 2.0, a gap extension penalty of 0.0, and an alignment score cut-off of 0.3. Sub-optimal alignments that meet the defined length and score thresholds are collected, along with the corresponding pLM-BLAST score for each alignment.
2. Transitivity has been shown to be a powerful approach in a number of sequence-related bioinformatics algorithms, such as MSA construction and homology search (36, 37). If residue *i* and residue *j* are aligned in one trace, and residue *j* and residue *k* are aligned in another trace, then residue *i* and residue *k* are assumed to be aligned in the third trace, called the transitive trace. For every two traces identified by pLM-BLAST, all possible transitive traces are generated and scored according to the cosine similarity substitution matrix derived from the initial pLM-BLAST self-alignment. To speed up the procedure and avoid introducing too much noise, only one round of transitivity is applied to suboptimal alignments.
3. After obtaining all the traces, the score matrix *Ms* is constructed by calculating the score of each cell *(i*,*j)* based on the collected traces *T*.

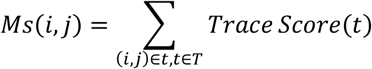
4. To estimate the repeat length, the scores of all cells located at the same distance (from 1 to *Lseq/2*) to the diagonal of the score matrix are summed separately for each distance, and all distances with a score greater than 0 are considered as possible lengths to be evaluated in the following steps.
5. For each potential repeat length *l* collected in the previous step, a sliding window of *l* residues is used to scan along the scoring matrix *Ms*. Following the procedure of HHrepID (16), we assume that conserved regions are more likely to be located in the middle of repeats, while the boundary regions are more prone to substitutions and indels. Considering this, we assign a weight to each column (residue) of a sliding window according to its position *p* to calculate the overall score. 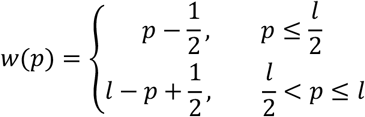 For each possible repeat length *l*, the position *p* of the representative repeat is determined by maximizing the total score in the region covered by a sliding window.

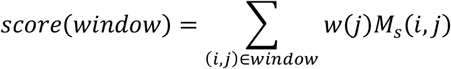
6. Once the position of the representative repeat is determined for a repeat length of *l*, its embedding is calculated by weighting the original repeat embedding according to the score matrix *Ms*.

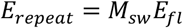

where *Msw* and *Efl* represent the columns of the score matrix *Ms* corresponding to the residue range of the representative repeat and full-length sequence embeddings, respectively. This scheme weights the embeddings of the corresponding sites using the columns of the score matrix, similar to creating a profile based on an MSA. In effect, the resulting weighted residue embeddings incorporate information collected in suboptimal alignments, thus improving the performance of the subsequent iterative repeat extraction step.
7. After obtaining the representative repeat embedding, another local pLM-BLAST comparison is performed to search it against the full-length sequence embedding to extract repeat instances. Each identified repeat is then compared to the representative repeat embedding using pLM-BLAST in global mode to derive the pairwise alignment and its score. These pairwise alignments are then combined to produce a multiple alignment. The results of different estimated repeat lengths are compared based on selected evaluation metrics (e.g. number of residues involved in all reported repeats) and the one with the best metrics is reported.
8. Once a search round is complete, the score matrix *Ms* is updated by masking all residue positions involved in the repeat ranges identified in that round. Steps 4-7 are then repeated with the updated score matrix until no more repeats are reported. An example of the analysis of domains containing more than one repeat type is shown in SI Fig. 1.

### Benchmark of protein repeat detection software

In addition to pLM-Repeat, two state-of-the-art self-alignment algorithms were selected for the performance benchmark: RADAR (v1.3, https://github.com/AndreasHeger/radar) (15), which operates on single sequences, and HHrepID (http://ftp.tuebingen.mpg.de/pub/protevo/HHrepID/) (16), which uses MSAs as input. Additionally, we evaluated the performance of HHrepID with a single sequence (no MSA) as input. Domains of both the positive and negative data sets were analyzed for each software. The benchmark was performed at two levels: protein level, and repeat level. In the protein level benchmark, a repeat protein was considered correctly detected by a given tool if more than half of its predicted repeats correctly aligned with at least one other predicted repeat. At the repeat level, each predicted repeat was considered correct if it correctly aligned with at least one ground truth repeat annotated in the RepeatsDB database. For non-repetitive domains identified as repeat proteins, we took the number of detected repeats as the number of false positives. In both benchmark modes, a correct alignment was defined as a structural alignment with a length-normalized TM-score greater than 0.5 and a sequence coverage greater than 50% provided by TM-Align (38). Repeat structures corresponding to detected residue ranges were extracted using the Biopython package (39), following the residue mapping dictionary provided in the localpdb package (40).

MSAs for all proteins in the repeat and non-repeat datasets were generated by searching each sequence against the UniRef30 database (version 2022-02) using HHblits with default settings (41). A series of repeat score thresholds, repeat family P-value thresholds, and self-sequence alignment score thresholds were evaluated for RADAR, HHrepID, and pLM-Repeat, respectively, to derive the benchmark results shown in Fig. 2. We also tested CE-symm (v2.2.2, https://github.com/rcsb/symmetry/tree/master) (18), a state-of-the-art structure-based repeat detection software, on the same datasets using default settings. To compare CE-symm with pLM-Repeat equipped with structure embeddings, the structures of the proteins included in the benchmark dataset were converted into embeddings by two inverse folding models, ESM-IF (42) and MIF (43), respectively. These embeddings were used as inputs to pLM-Repeat with the transitivity turned off to avoid introducing noise and other settings left as defaults.

**Figure 2.**
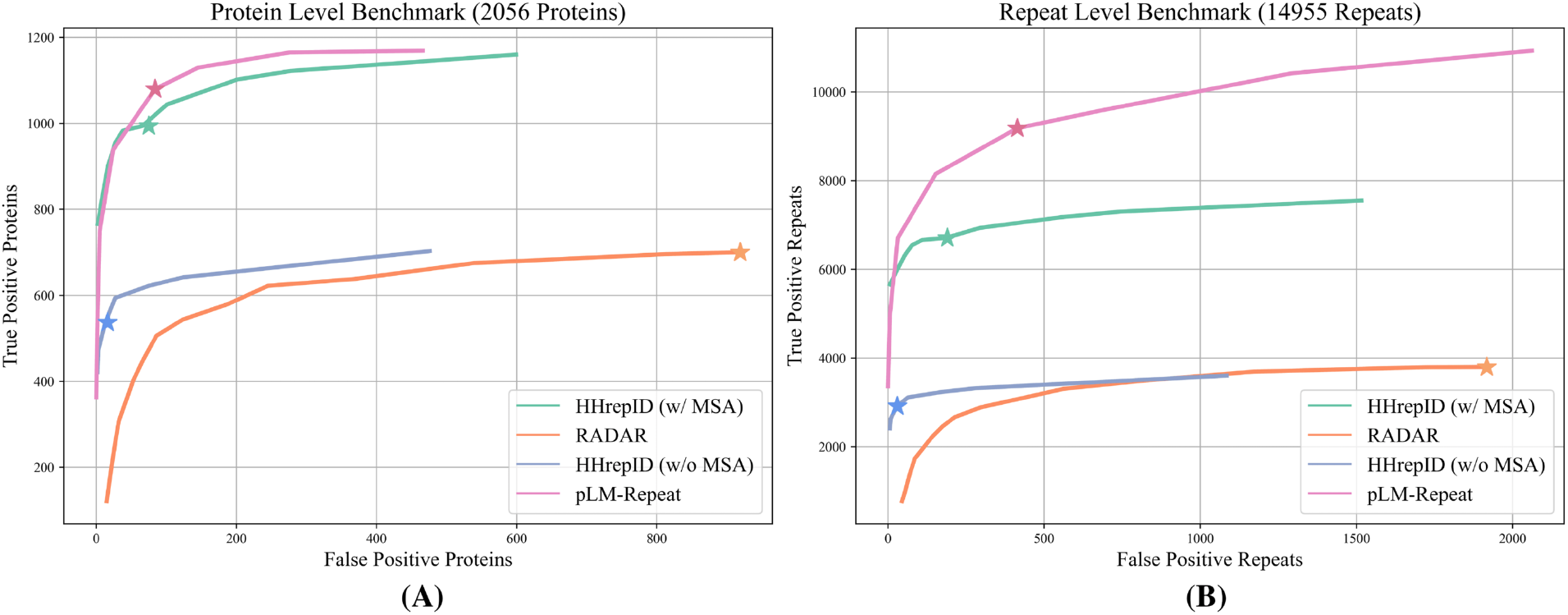
Benchmark results. Performance benchmark of sequence-based repeat detection methods. Repeat family P-value threshold, repeat score threshold, and self-sequence alignment score threshold are tuned for HHrepID (with or without MSA), RADAR, and pLM-Repeat, respectively. The performance of each software at default settings is marked with an asterisk. Benchmarking was performed at the protein (A) and repeat (B) level (see Methods for details).

We performed the speed comparison by analyzing proteins of different lengths using RADAR, HHrepID and pLM-Repeat. The speed test was performed on the same system equipped with an AMD EPYC 7742 64-core CPU. The runtime for each query in each software was averaged in three replicates.

### Implementation of DeepRepeat

To enable fast pre-filtering in large scans, we trained a neural network called DeepRepeat to identify repeats with patterns similar to those found in known repeat proteins (Fig. 5A). DeepRepeat uses slightly modified light attention architecture proposed by Stärk et al. (44). The model takes as input a per-residue embedding, which is also the input of pLM-BLAST. The embedding is transformed by two separate 1D convolution layers, both with a filter size of 9 and an output channel of 1024. The output of the first convolution layer is followed by a `Softmax` layer to generate attention distributions, while the output of the second convolution layer is followed by a `Dropout` layer and results in feature maps. The Hadamard product of the attention distributions and feature maps is summed along the sequence length dimension to generate the weighted sum, which is then concatenated with the feature maps after the `MaxPool` operation. The resulting fixed shape embeddings of the input samples are then fed into a linear layer for binary classification. The model was trained using the Adam optimizer at a learning rate of 1×10^-6^, and the dataset was split into training and validation sets at a ratio of 9:1 for the cross-validation training routine. Binary cross-entropy loss was used for training. Precision, recall and F1 score were evaluated to assess the performance of the model.

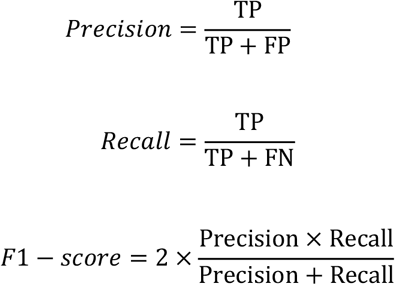

**Figure 5.**
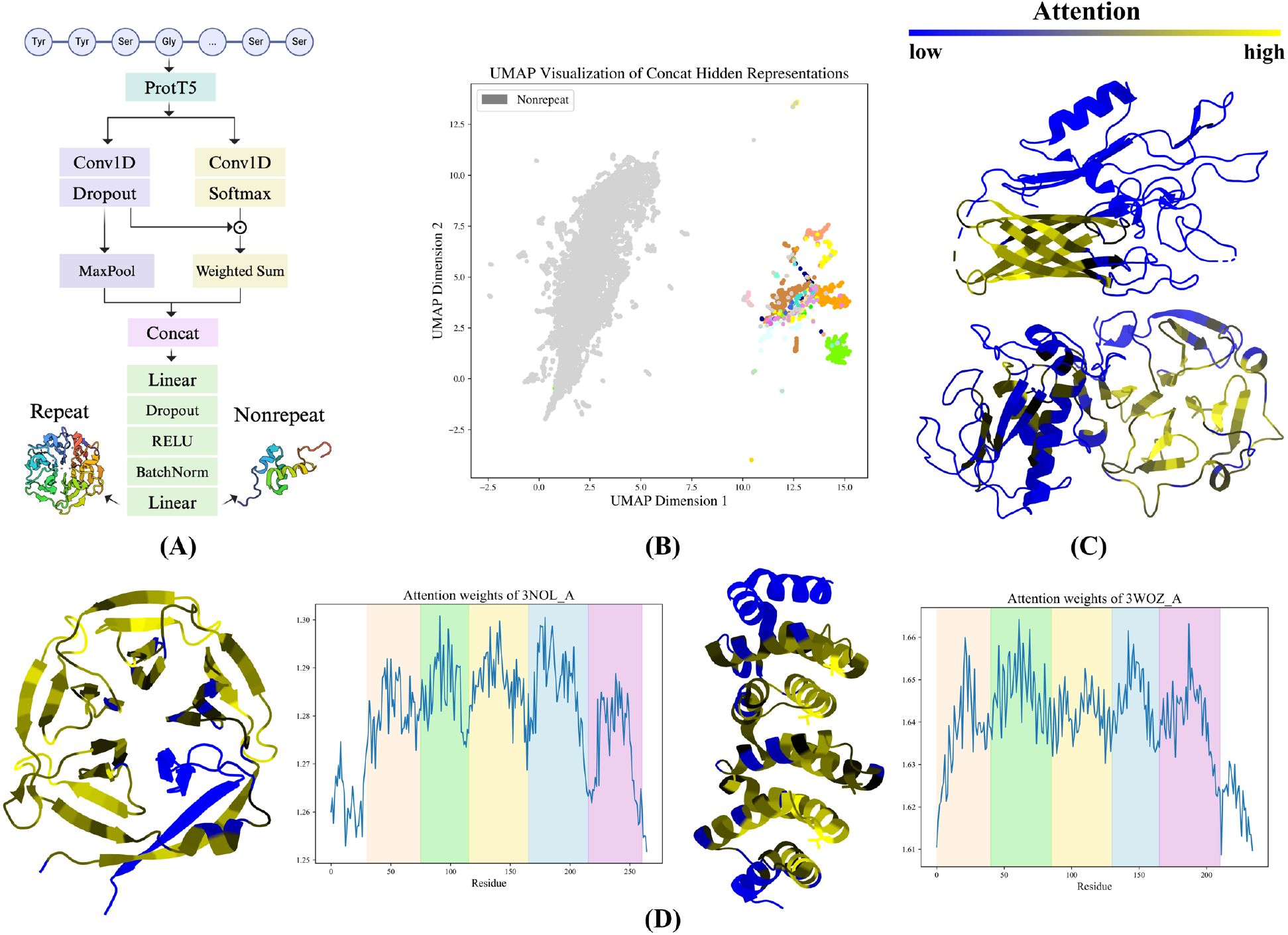
A neural network for distinguishing between known repetitive and non-repetitive proteins. (A) The model architecture, including a light attention module and a prediction layer. (B) The UMAP visualization of all proteins with and without repeats in the compiled dataset. Nodes of non-repeat proteins are colored in gray, while nodes of repeat proteins are colored in different colors based on fold class. (C) Protein structures colored based on attention scores, with blue and yellow representing low and high attention scores, respectively. Repeat domains in 3UAQ_A (top) and 1FBL_A (bottom) show higher attention scores than their non-repeat counterparts. (D) Repeat proteins 3NOL A (left) and 3WOZ A (right) show periodic attention patterns.

To explore the potential interpretability of the neural network, raw attention weights with a shape (*Lseq*, 1024) were extracted for each protein of length *Lseq* before applying the `Softmax` layer. The average across the 1024 dimensions was assigned as the attention score for each residue and mapped to protein structures for visualization. We also examined the embeddings of all repeat and non-repeat proteins obtained from the `Concat` layer, which is the last layer of the light attention module. Each embedding from the `Concat` layer has a fixed size (2048) regardless of sequence length. All protein embeddings were projected into 2D space and visualized by the Uniform Manifold Approximation and Projection (UMAP) framework using the UMAP package (45). The model was implemented in the PyTorch framework (v1.13.1).

### Scan on the AFDB90v4 database

The AFDB90v4 database was downloaded from the Uniprot3D website (https://uniprot3d.org/) (46). Each protein in this database is assigned a “functional brightness”, defined as the coverage of the sequence with annotations of homologs, ranging from 0 to 1. 682,563 sequences with low functional brightness (≦ 0.1) and no representatives were collected from the AFDB90v4 database, converted to embeddings using the ProtT5 model, and passed to the DeepRepeat model. Entries recognized as repeat proteins were further clustered to 50% sequence identity and 80% coverage using MMseqs2. We assessed their sequence novelty by searching them against the ECOD70 database (47) using HHSearch (48) with default settings and discarding those reporting hits with a probability greater than 30%. The predicted structures of the remaining proteins were retrieved using the AlphaFold database API and searched against the PDB100 database using Foldseek (49) with an E-value threshold of 0.1 to identify structures that had no similar counterparts in the PDB. These filters resulted in a data set of 5460 proteins of potential interest (S2 File).

### Analysis of selected repeat folds

We examined several selected repetitive folds using a variety of tools: homology search using HHpred and pLM-BLAST, repeat detection using HHrepID and pLM-Repeat, structure search using Foldseek, and sequence clustering using MMseqs2. The majority of these analyses, with the exception of our newly developed tools and Foldseek, were performed using the MPI bioinformatics toolkit (https://toolkit.tuebingen.mpg.de/) (50).

### Visualization

Structures were visualized with PyMOL3.0. Figures 1 and 5A were created with BioRender.com. Other figures were plotted using the Matplotlib and seaborn packages.

## Results

### pLM-Repeat benchmark

pLM-Repeat identifies repeat patterns in a given sequence based on suboptimal self-alignments obtained with pLM-BLAST (see Methods for details). We evaluated the performance of pLM-Repeat on the compiled set of 2056 repeat and 1987 non-repeat protein domains along with two sequence-based self-alignment methods, HHrepID and RADAR. In contrast to RADAR, which works with a single sequence input, HHrepID can take either a single sequence as input or an MSA derived from that sequence, the latter offering significantly better accuracy at the expense of speed. The benchmark procedure includes pLM-Repeat, RADAR and HHrepID in two modes and was performed in two variants, first to evaluate the ability to discriminate between repetitive and non-repetitive proteins and second to evaluate the accuracy of predicting individual repeats.

The results of the first benchmark were plotted as the true positives vs. false positives plot at different significance thresholds (Fig. 2A). As expected, both pLM-Repeat and standard HHrepID significantly outperformed the single sequence-based methods RADAR and HHrepID without MSAs, identifying approximately twice as many repeat proteins with a false positive rate of 10%. With default settings, pLM-Repeat showed comparable performance to HHrepID, identifying 1080 and 994 correct repeat domains, respectively, with a similar number of false positives of 84 and 66 incorrect domains, respectively. Notably, HHrepID can still detect 766 repeat proteins without false positives at a very stringent P-value threshold of 1×e^−13^, while 363 repeat domains can be identified without false positives at a score threshold of 0.5, indicating better false positive control due to the statistical evaluation framework used in HHrepID.

All correct predictions from the first benchmark were obtained at a self-alignment score threshold of 0.3, a repeat family P-value threshold of 0.001, and a repeat score threshold of 20 for pLM-Repeat, HHrepID, and RADAR, respectively, and were further categorized based on the RepeatsDB fold classes as shown in Table 1. Although our work focuses on sequence-based repeat detection, we also included predictions from a structure-based software, CE-symm. This method was originally designed to detect structural repeats in the β-propeller and TIM-barrel domains, and thus provides a reference point for assessing potential bias in the evaluated methods. Among all sequence-based methods included in our benchmark, HHrepID and pLM-Repeat consistently showed top performance, nearly dominating each fold class. pLM-Repeat showed a bias towards certain protein folds, such as β-propellers, where it correctly identified 259 (70.0%) domains, approaching the performance of the structure-based method CE-symm. On the other hand, HHrepID outperformed other methods in the detection of β-solenoid and α/β-solenoid domains, with 101 (56.1%) and 112 (74.7%) correct detections, respectively, outperforming even CE-symm. Interestingly, pLM-Repeat performed best in two repeat protein folds with highly diverse repeat sequences but obvious structural repeat patterns, β-barrels/hairpins and α-solenoids, detecting 41 (17.9%) and 336 (73.5%) domains, respectively, compared to 18 (7.9%) and 269 (58.9%) by the second-best method, HHrepID. SI Fig. 2 and SI Fig. 3 show a selection of correctly identified domains of both folds by pLM-Repeat. Furthermore, lowering the pLM-Repeat score threshold to 0.25 (corresponding to the increase in false positive rate from 4.2% to 13.9%) resulted in 85 more correctly predicted proteins, mostly TIM-barrels, increasing from 45 (12.2%) to 107 (29.0%; SI Table 1). Visualization of the detected domains shows that pLM-Repeat successfully identified the β-α repeat units in most TIM barrels, despite the low internal repeat identity (SI Fig. 4). Such apparent structural periodicity, combined with low repeat sequence identity in protein folds like β-barrels and TIM-barrels, introduces ambiguity in determining whether these structural repeats result from true duplication events and poses a challenge to sequence-based detection approaches. The promising performance of pLM-Repeat on these difficult targets underscores its potential for evolutionary analysis, since only sequence similarity, as opposed to structural similarity, reliably indicates potential homology between repeat units.

**Table 1.** Number of correctly detected proteins on RepeatsDB dataset based on fold classes.

| Repeat Protein Fold | pLM-Repeat | HHrepID | HHrepID-single | RADAR | CE-symm |
| --- | --- | --- | --- | --- | --- |
| β-Solenoid (180) | 71 (39.4%) | **101** (56.1%) | 43 (23.9%) | 62 (34.4%) | 80 (44.4%) |
| $\alpha/\beta$ Solenoid (150) | 104 (69.3%) | **112** (74.7%) | 100 (66.7%) | 60 (40.0%) | 106 (70.1%) |
| α-Solenoid (457) | **336** (73.5%) | 269 (58.9%) | 145 (31.7%) | 217 (47.5%) | 306 (67.0%) |
| $\beta$ Hairpins (40) | 17 (42.5%) | **25** (62.5%) | 14 (35.0%) | 13 (32.5%) | 17 (42.5%) |
| Box (57) | 40 (70.2%) | **44** (77.2%) | 0 (0.0%) | 14 (24.6%) | 44 (77.2%) |
| TIM-Barrel (370) | 45 (12.2%) | 28 (7.6%) | 8 (2.1%) | **57** (15.4%) | 101 (27.3%) |
| β-barrel/hairpins (229) | **41** (17.9%) | 18 (7.9%) | 1 (0.4%) | 9 (3.9%) | 54 (23.6%) |
| β-Propeller (370) | **259** (70.0%) | 216 (58.4%) | 126 (34.1%) | 141 (38.1%) | 261 (70.5%) |
| $\alpha/\beta$ Prism (25) | 22 (88.0%) | **23** (92.0%) | 2 (8.0%) | 16 (64.0%) | 25 (100.0%) |
| α-Barrel (28) | **24** (85.7%) | 21 (75.0%) | 9 (32.1%) | 0 (0.0%) | 22 (78.6%) |
| $\alpha/\beta$ Trefoil (49) | 38 (77.6%) | **39** (79.6%) | 18 (36.7%) | 22 (44.9%) | 31 (63.3%) |
| Aligned prism (15) | 7 (46.7%) | **10** (66.7%) | 1 (6.7%) | 4 (26.7%) | 15 (100.0%) |
| α-Beads (7) | **7** (100.0%) | 6 (85.7%) | 5 (71.4%) | 5 (71.4%) | 7 (100.0%) |
| β-Beads (23) | **18** (78.3%) | 17 (73.9%) | **18** (78.3%) | **18** (78.3%) | 6 (26.1%) |
| $\alpha/\beta$-Beads (26) | 18 (69.2%) | **23** (88.5%) | 20 (76.9%) | 19 (73.0%) | 13 (50.0%) |
| $\beta$ Sandwich beads (21) | **16** (76.2%) | 15 (71.4%) | 11 (52.4%) | 9 (42.9%) | 8 (38.1%) |
| $\alpha/\beta$ Sandwich (32) | 12 (37.5%) | **18** (56.3%) | 12 (37.5%) | 13 (40.6%) | 6 (18.8%) |

All software was used with default settings to obtain the statistics for each fold class (with the number of domains in parentheses) in this table. Statistics for the method that achieved the best performance among all sequence-based approaches are highlighted in bold. CE-symm performance is included for ease of comparison.

Figure 2B shows the results of the second benchmark variant, which focuses on the detection of individual predicted repeats by comparing them to those deposited in RepeatsDB. Again, both pLM-Repeat and HHrepID report more than twice as many correct repeats as RADAR and HHrepID without MSAs. In addition, HHrepID again shows exceptional accuracy with a low false positive rate, achieving up to 6710 correct repeats versus only 190 incorrect repeats with default settings. A notable difference in this benchmark is that pLM-Repeat detects a much larger number of repeats than HHrepID. For example, pLM-Repeat identifies 9182 repeats with a default SSA-score threshold of 0.3, while also reporting 414 false positives. This discrepancy occurs because pLM-BLAST excels at detecting short but significant local alignments and therefore includes a wider range of repeats within domains.

Encouraged by the above results showing the potential of protein sequence representations in repeat detection, one might wonder whether protein structure embeddings could be integrated into our pLM-Repeat workflow. Since pLM-BLAST was designed as a universal tool that can be combined with any embedder, we explored this prospect by feeding pLM-Repeat with structure embeddings derived from two inverse folding models, ESM-IF (42) and MIF (43). pLM-Repeat was run with default settings, except for turning off transitivity to reduce unnecessary noise. Despite some successful cases where pLM-Repeat reported correct repeats from structure embeddings (SI Fig. 5), the overall benchmark on the RepeatsDB dataset showed that CE-symm significantly outperformed pLM-Repeat with structure embeddings (SI Fig. 6), while MIF representations achieved slightly more correct detections than ESM-IF representations, with 305 and 245 domains, respectively. This inferior performance may be due to the fact that the feasibility of comparing embeddings to detect structurally similar segments has not been validated, and pLM-BLAST, although technically accepting any embeddings, was mostly tested with the ProtT5 model.

Speed is another important feature of sequence analysis software, especially for large scans. We compared the run times of the software included in the performance benchmark using proteins of different lengths (see Table 2). RADAR demonstrated rapid detection within 0.1 seconds for all sequences, followed by HHrepID without MSA, which completed the run within seconds. In contrast, the standard HHrepID, including an HHblits search against the UniRef30 database, took minutes to complete each job, while pLM-Repeat performed analyses significantly faster than HHrepID. For example, pLM-Repeat and HHrepID took 24.7 and 272.2 seconds, respectively, on the domain 2QJ6_A.

**Table 2.** Speed comparison of sequence-based repeat detection methods.

| Query (Length) | pLM-Repeat | HHrepID | HHrepID-single | RADAR |
| --- | --- | --- | --- | --- |
| 6A57_A (140) | 7.4s | 285.3s | 1.2s | 0.1s |
| 4DB6_A (211) | 9.4s | 271.5s | 1.8s | 0.1s |
| 2QJ6_A (332) | 24.7s | 272.2s | 2.4s | 0.1s |
| 5AMS_A (431) | 23.1s | 606.7s | 0.6s | 0.1s |

All software was used with default settings to obtain the statistics in this table. The runtimes of pLM-Repeat and HHrepID include the step of obtaining protein sequence embeddings and multiple sequence alignments, respectively. HHrepID on a single sequence took only 0.6 s to complete the procedure on the 5AMS\_A domain because it detected a few suboptimal alignments.

### Examples of pLM-Repeat performance

In this section, we illustrate the performance of pLM-Repeat with examples of different repeat folds. Chain A of protein PDB:2X19 is an Armadillo repeat protein containing 22 repeats with a low repeat sequence identity of 16.6% on average (Fig. 3A). pLM-Repeat successfully recognized 17 repeats, most of which were α-hairpins, while HHrepID with 3 rounds of the HHblits search to generate MSAs failed to report any repeats. We also applied pLM-Repeat to a pectin lyase-like β-solenoid domain (PDB: 7C7D) (Fig. 3B). Despite the low average internal sequence identity of 16.6%, 11 out of 12 repeats were correctly detected, while HHrepID identified 10 incomplete repeats. In a 22-stranded β-barrel with an average repeat identity of 17.0%, pLM-Repeat reported 12 repeat fragments, of which 8 were correctly superimposed as β-hairpins (Fig. 3C). In the same sequence HHrepID recognized 5 repeat instances composed of incomplete β-hairpins. Finally, analysis of a tryptophan synthase TIM barrel revealed 7 β-α fragments with an average sequence identity of 17.8% (Fig. 3D), of which no repeats were identified by HHrepID.

**Figure 3.**
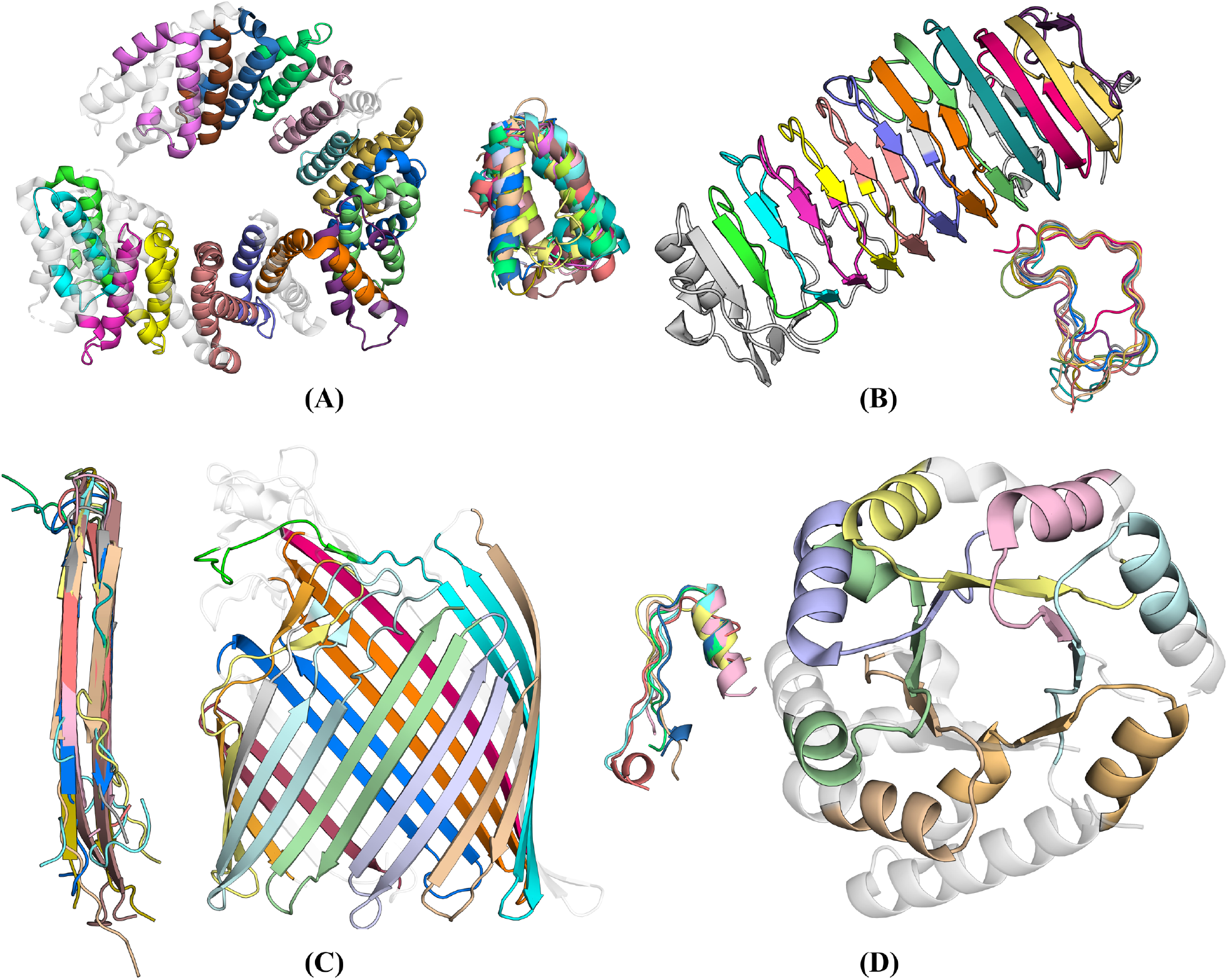
Performance cases of pLM-Repeat. Repeat detection results for (A) 2X19_B (repeat identity 16.6%), (B) 7C7D_A (16.6%), (C) 5NEC_A (17.0%), and (D) 3NAV_A (17.8%). Structures are colored according to the identified repeat regions, with repeats superposed to show structural similarity. Residues not included in the detected regions are shown in white with semi-transparency.

Another example of pLM-Repeat outperforming other methods in difficult cases is chain A of protein PDB:1W6S (Fig. 4). This domain consists of an 8-blade quinoprotein alcohol dehydrogenase-like propeller containing highly divergent repeats (internal repeat sequence identity of 20.8%) with a significant number of indels. RADAR struggled with this domain due to the interference caused by these massive indels, resulting in structurally dissimilar and incorrect repeats (Fig. 4D). Although HHrepID performed better by returning four repeats (Fig. 4C), only two of them corresponded to a 4-stranded β-meander propeller blade, while the rest were reducible repeats containing two blades (shown in green and yellow). The score matrix generated by pLM-Repeat clearly showed internal local alignment traces along a sliding window (Fig. 4A), and seven structurally similar repeats were successfully reported (Fig. 4B). It should be noted, however, that pLM-Repeat missed complete blades as repeats and only detected the β3-β4 hairpin of each β-meander. This limitation is due to massive insertions connecting the loop between β2 and β3 in some blades. Nevertheless, in this case, only pLM-Repeat provided a result in which the predicted repeat units are evolutionarily meaningful and structurally alignable.

**Figure 4.**
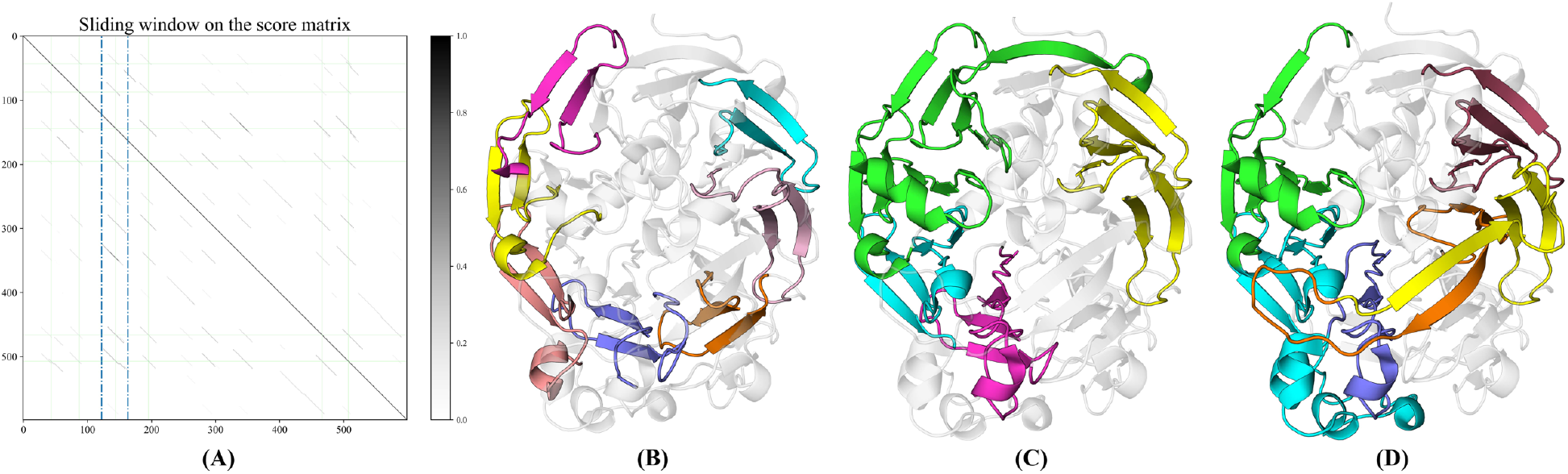
Performance comparison. Performance comparison between pLM-Repeat (B), HHrepID (C), and RADAR (D) on case 1W6S. A: Score matrix with the determined sliding window. Eight traces are clearly shown within the region of the sliding window. Green lines in the score matrix indicate positions of conserved Trp located at the fourth β-strand from 6 propeller blades.

### Distinguish between proteins with and without repeats using a deep learning model

While speed improvements have been achieved with pLM-Repeat, it is not efficient enough to perform large-scale scans on ultra-large sequence databases such as UniProt. To overcome this limitation, an auxiliary pre-filtering model, DeepRepeat, was developed to speed up such scans (Fig. 5; see Methods for details). The model is based on a neural network and was trained to detect whether input sequences contain patterns typical of known repeat proteins, using the same embeddings that serve as input to pLM-Repeat. Visualization of the internal representations obtained from the trained DeepRepeat network shows a clear separation between repeat and non-repeat proteins, which is critical for achieving accurate binary classification. In fact, the model achieved an F-score of 0.91 on the independent validation set, demonstrating its effectiveness in detecting repeat proteins. Since the model incorporates a light attention module (44), it is possible to extract its attention scores and indicate which regions of a given protein were essential for the prediction. For proteins containing both repetitive and non-repetitive regions, the former had significantly higher attention scores. This not only indicates that the model learned to detect relevant signals associated with repetition, but also suggests its potential to localize the regions containing repeats. Finally, we noticed that the attention weights on some repeat regions already showed a clear periodicity pattern, as shown in Figure 5D, further confirming the high performance of the trained model.

### Gallery of AFDB90v4 repeat proteins

To demonstrate the practical application of our developed tools in conjunction with the ever-increasing amount of protein sequence data, we performed a large-scale scan on the AFDB90v4 database (46), which was generated by clustering to a maximum sequence identity of 50% of the UniRef database and filtered based on a pLDDT threshold of 90 to retain only proteins with confidently predicted models. Each sequence within AFDB90v4 is assigned a “functional brightness” index, ranging from 0 to 1 according to the annotation coverage provided in different databases. We focused on entries with low functional brightness scores, as these unknown proteins are more likely to exemplify undefined families. The embeddings of 682,563 proteins were first fed into the into DeepRepeat model, and those predicted to contain repeats were subjected to further analysis to evaluate repeat patterns and assess sequence and structure novelty using pLM-Repeat and other tools (see Methods for details).

Although the DeepRepeat neural network was trained to detect repeat protein domains based on prior knowledge rather than *de novo* repeat patterns, some of the proteins predicted to contain repeats exhibit sequence and structure novelty relative to defined repeat folds. Figure 6 shows a selection of potentially novel repeat proteins detected by our pipeline. One protein of significant interest (UniProt ID: A0A7C3HQW7) contains repeats that fold into twisted long β-hairpins, deviating from typical β-barrel hairpins (first structure in the third row in Fig. 6, see also SI Fig. 7A). Interestingly, although the inner core of this structure resembles a β-barrel, no homology to any β-barrel could be detected using HHpred or pLM-BLAST searches, regardless of whether the search was performed with the full-length sequence or only the barrel region. Some of the close homologs of this protein show structural diversity, such as the one with UniProt ID A0A534S4A8, which contains three repeats instead of four (SI Fig. 7B). Another protein with a novel repeat topology is A0A424SVE7. Its repeat unit starts with a small β-hairpin, followed by an α-helix and an elongated β-hairpin composed of several separate β-strands that spiral up and down (second structure in the third row in Fig. 6, see also SI Fig. 8). Another structurally novel fold is A0A0S8GK70, which is characterized by 5-fold internal symmetry and repeats with three β-strands, the first and last of which interact with adjacent repeats (second structure in the second row in Fig. 6, see also SI Fig. 9).

**Figure 6.**
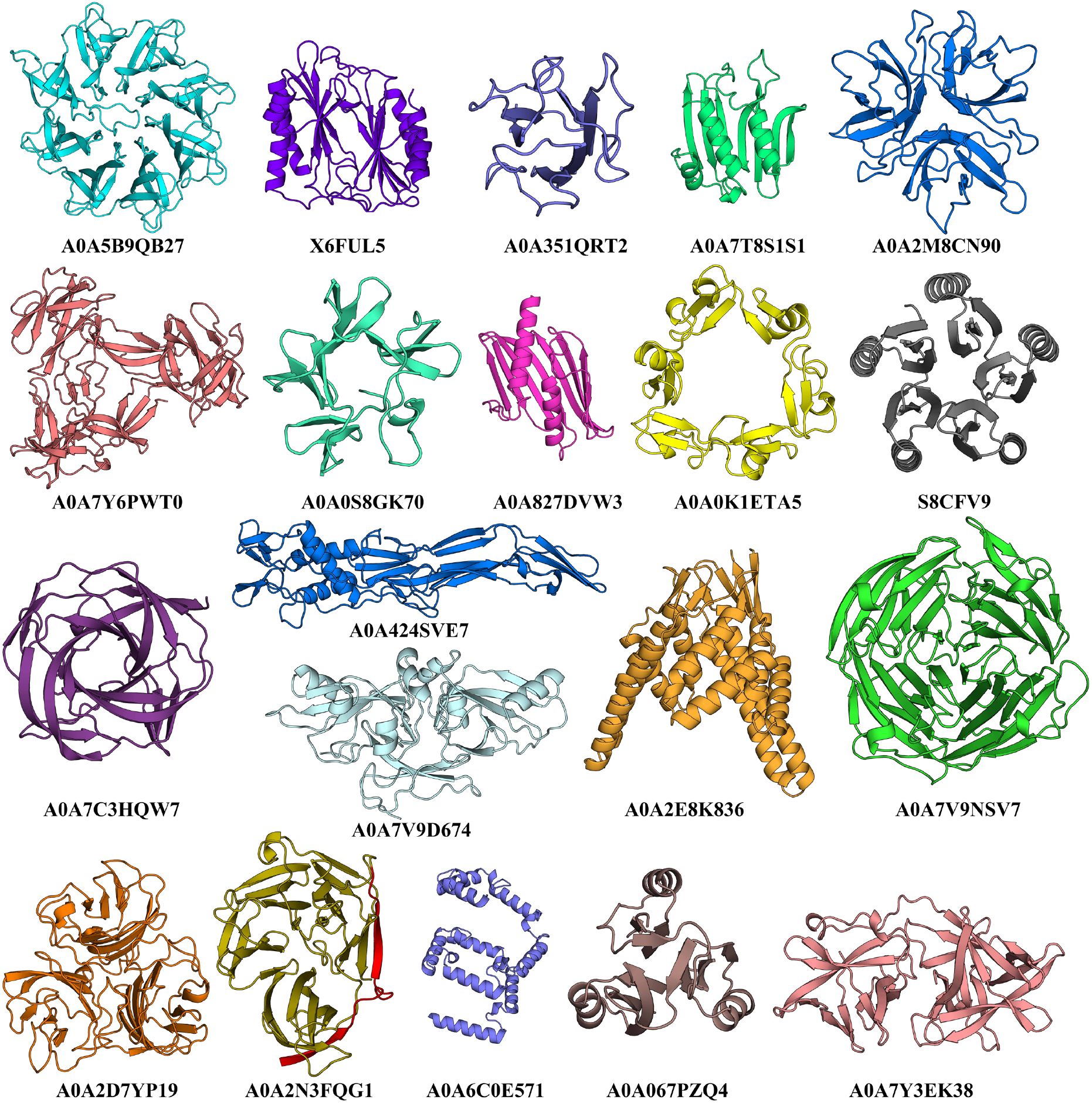
Gallery of potentially novel repeat proteins. A collection of potentially novel repeat proteins detected in the AFDB90v4.

## Discussion

Sequence representations derived from protein language models (pLMs) provide new opportunities to address classical bioinformatics tasks related to proteins, such as homology detection, classification, design, and function annotation. In this study, we investigated the application of protein sequence embeddings for repeat pattern detection. The performance of self-comparison algorithms for repeat identification is highly dependent on the quality of the collected suboptimal alignments. By exploiting the sensitivity observed in remote homology detection with pLM-BLAST, we are able to generate a set of internal repeat alignments at the initial stage of the workflow, which forms the cornerstone of pLM-Repeat’s performance, further strengthened by the strategy of transitivity and profile-like repeat embedding. pLM-Repeat shows promising repeat detection performance comparable to well-established methods such as HHrepID. While HHrepID enhances self-alignment signals by HMM-HMM comparison, pLM-Repeat is characterized by its independence from time-consuming MSA construction, as it uses enriched representations obtained from pLMs trained on millions of protein sequences.

The performance of pLM-Repeat in discriminating between repeat and non-repeat proteins is comparable to that of HHrepID, with faster run times and detection of more repeat units (Fig. 2). We found that pLM-Repeat achieved the most correct detections in several specific repeat folds, such as β-propellers, α-solenoids, and β-barrels. In addition, a slight decrease in the self-alignment score threshold allows for many more correct detections in TIM-barrels. This may be due to pLM-BLAST’s capability to detect short locally similar fragments, especially since repeat units in these protein families are relatively short (20-40 residues), making them better covered by pLM-Repeat and suggesting the complementary role of the pLM-Repeat workflow to HHrepID. To enable the application of pLM-Repeat to large databases, we trained DeepRepeat, a deep learning model that uses a lightweight attention module, an architecture that has been shown to be effective in various transfer learning tasks, such as predicting protein localization (44) and the optimal pH of enzymes (51). This model serves as a fast prefilter to identify proteins that have patterns similar to well-characterized repeat folds based on prior knowledge.

The ever-increasing amount of sequence and structure data represents a valuable resource for various purposes, such as exploring statistical trends in protein evolution (52) or identifying structurally novel domains (53). Many efforts have focused on annotating the AlphaFold database (AFDB) to improve its accessibility to the community (46, 54, 55). For example, a recent study created a comprehensive domain encyclopedia based on the entire AFDB by applying domain segmentation approaches and reported symmetrical domains defined according to the structural symmetry Z-score calculated by SymD (55). In this work, we used DeepRepeat and pLM-Repeat to scan protein sequences with low “functional brightness” annotated in AFDB90v4 (46), as these proteins have predicted structures in AFDB for easy manual verification and validation if the detected protein indeed contains a repeat. This pipeline led to the discovery of a set of repeat domains exhibiting sequence and structural novelty, some of which were particularly intriguing, such as the 4-copy β-barrel-like domain composed of twisted β-hairpins. More rigorous assessment of novelty and further classification of these repeat domains will be essential to advance repeat protein studies.

A limitation of pLM-Repeat is the lack of rigorous statistical evaluation, beyond from the alignment score provided by pLM-BLAST. Traditional statistical frameworks, such as the extreme value distribution theory used in HHrepID, may not be appropriate for protein embeddings because shuffling residue embeddings would disrupt the context dependency of pLM representations. The lack of a robust statistical framework may challenge to the pLM-Repeat procedure in certain steps. For instance, one observed problem is the redundancy of detected traces that are only a few residues apart from each other. While HHrepID severely penalizes such shifted alignments, they may still pass the subpath selection step if they exceed the given threshold. One solution is to increase the default Sigma factor in pLM-BLAST. The default Sigma factor of 2.0 means that subpaths reported by pLM-BLAST are limited to a minimum score of two times the average standard deviation of the substitution matrix, so increasing it will make the subpath selection more stringent. Alternatively, scanning the representative embeddings along the full-length sequence embeddings with different pLM-BLAST score thresholds and reporting the best result can avoid missing repeat proteins with such behavior (see SI Fig. 10 for details on the importance of the Sigma factor parameter for prediction).

Another shortcoming is the output of multiple alignments of the detected repeat units. Currently, pLM-Repeat generates the alignment of all detected repeats by simply concatenating pairwise alignments of each detected repeat to the reference repeat (Fig. 1), resulting in alignments that are not optimal. Recent work using pLM embeddings for MSA construction may provide a possible solution. However, most of the currently used strategies, such as vcMSA (27) and PEbA (56), rely on clustering and ordering of residue embeddings and are designed to process gapless sequences as input. Consequently, they fail to take advantage of the valuable pairwise alignment obtained during the self-alignment process.

Finally, we envision further enhancements to the DeepRepeat model that would enable it to detect *de novo* repeat patterns encoded in sequence embeddings and even report repeat regions. This capability has been implemented at the structural level in DeepSymmetry, which detects structural repeats and density maps using 3D convolutional networks (57), suggesting that such a generalization of the DeepRepeat model is feasible.

In summary, we have explored the use of protein sequence embeddings for repeat identification using pLM-Repeat, a repeat pattern detection pipeline, and DeepRepeat, a network trained to predict repeat domains according to prior knowledge. Our tools achieve promising sensitivity as well as good speed, and excel in some specific protein folds, offering a complementary approach to existing methods such as HHrepID, which relies on MSAs, especially in large-scale analysis, where we present a set of potentially novel repeat domains from the AFDB90v4 scan.

## Acknowledgements

We would like to thank Wenfei Xian (Max Plank Institute for Biology) and Yandong Wen (Max Planck Institute for Intelligent Systems) for insightful discussions and Kamil Kaminski (University of Warsaw) for help with the usage of pLM-BLAST. Computation was performed on the MPI-BIO cluster and the HPC system Raven at the Max Planck Computing and Data Facility. K.Q. would like to thank the IMPRS From Molecules to Organisms PhD program. This work was supported by institutional funds of the Max Planck Society.

## Data availability

Source codes of pLM-Repeat, together with trained model weight and generated datasets are available on the GitHub repository https://github.com/KYQiu21/plmrepeat.

