## Supplementary File 1 for "Exploiting protein language model sequence representations for repeat detection"

### Supplementary Figures

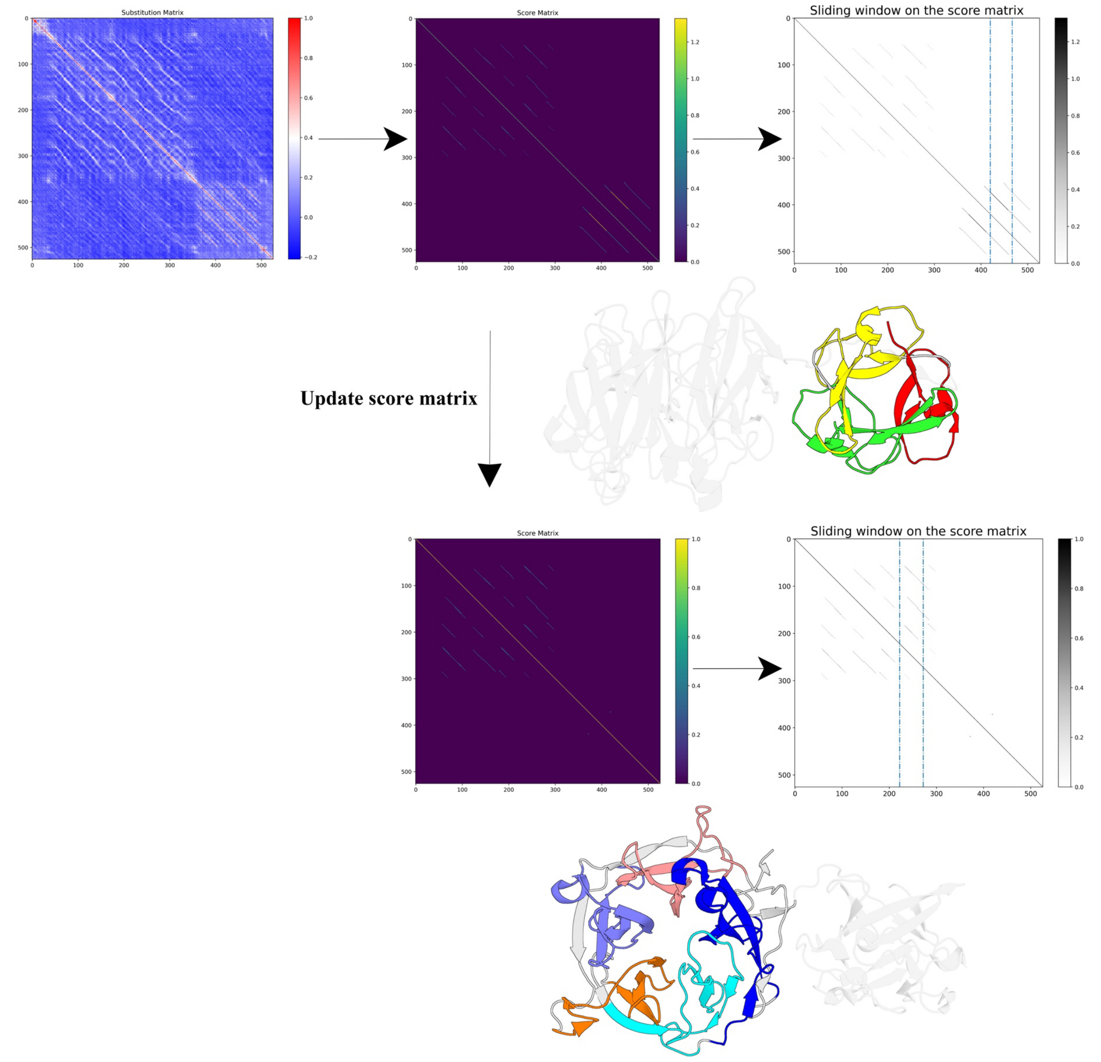


**Supplementary Figure 1:** pLM-Repeat analysis on the domain 3VT1_A, which contains two types of repeats. 5 propeller repeats and 3 trefoil repeats were reported.

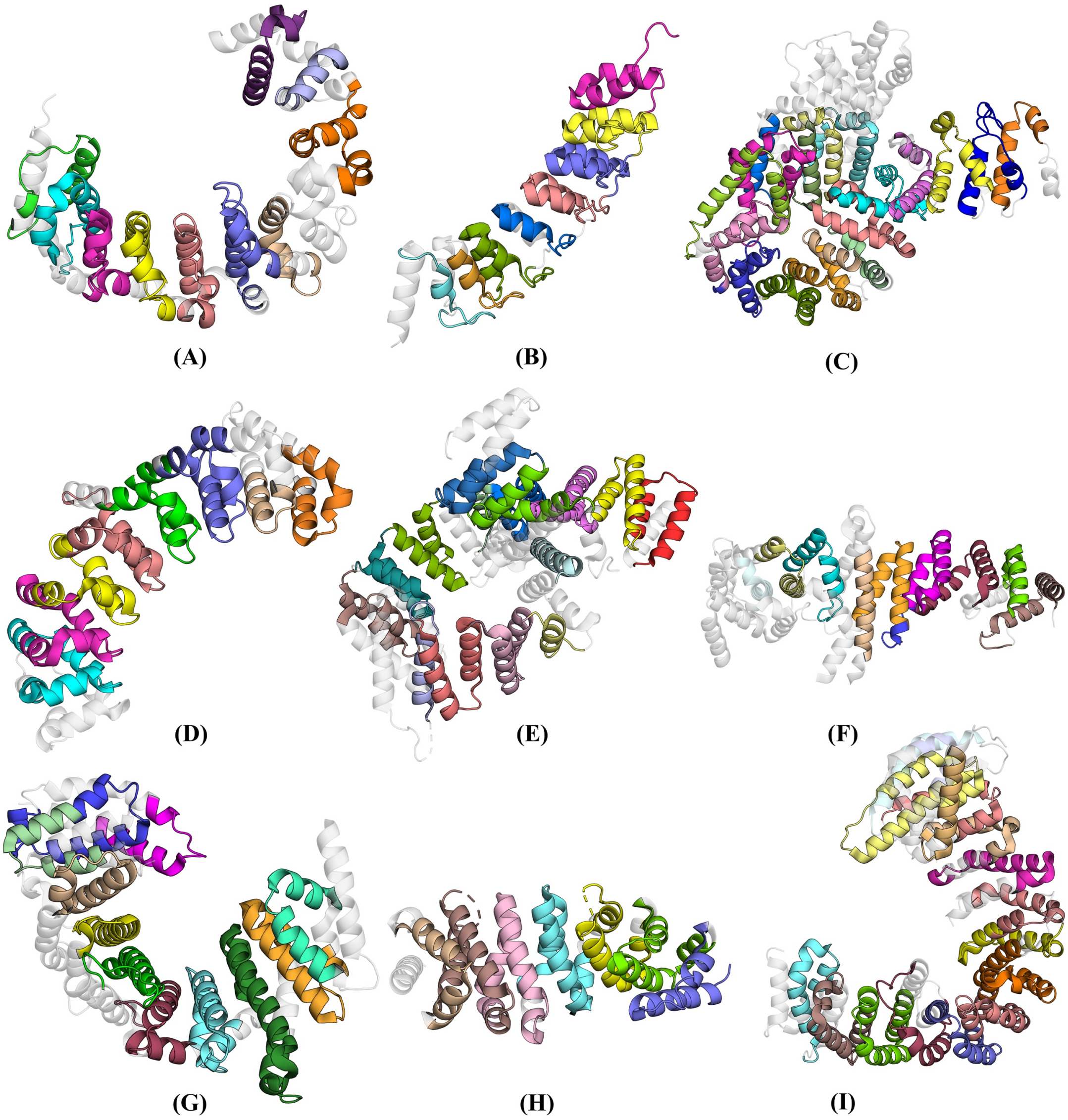


**Supplementary Figure 2:** A selection of α-solenoid domains correctly detected by the pLM-Repeat method. Protein structures are colored according to repeat ranges. (A) 4A3V_A, (B) 6BY9_A, (C) 6QDK_A, (D) 3V71_A, (E) 5NNP_A, (F) 3TJ1_A, (G) 4GMO_A, (H) 3T7U_A, and (I) 2QNA_A.

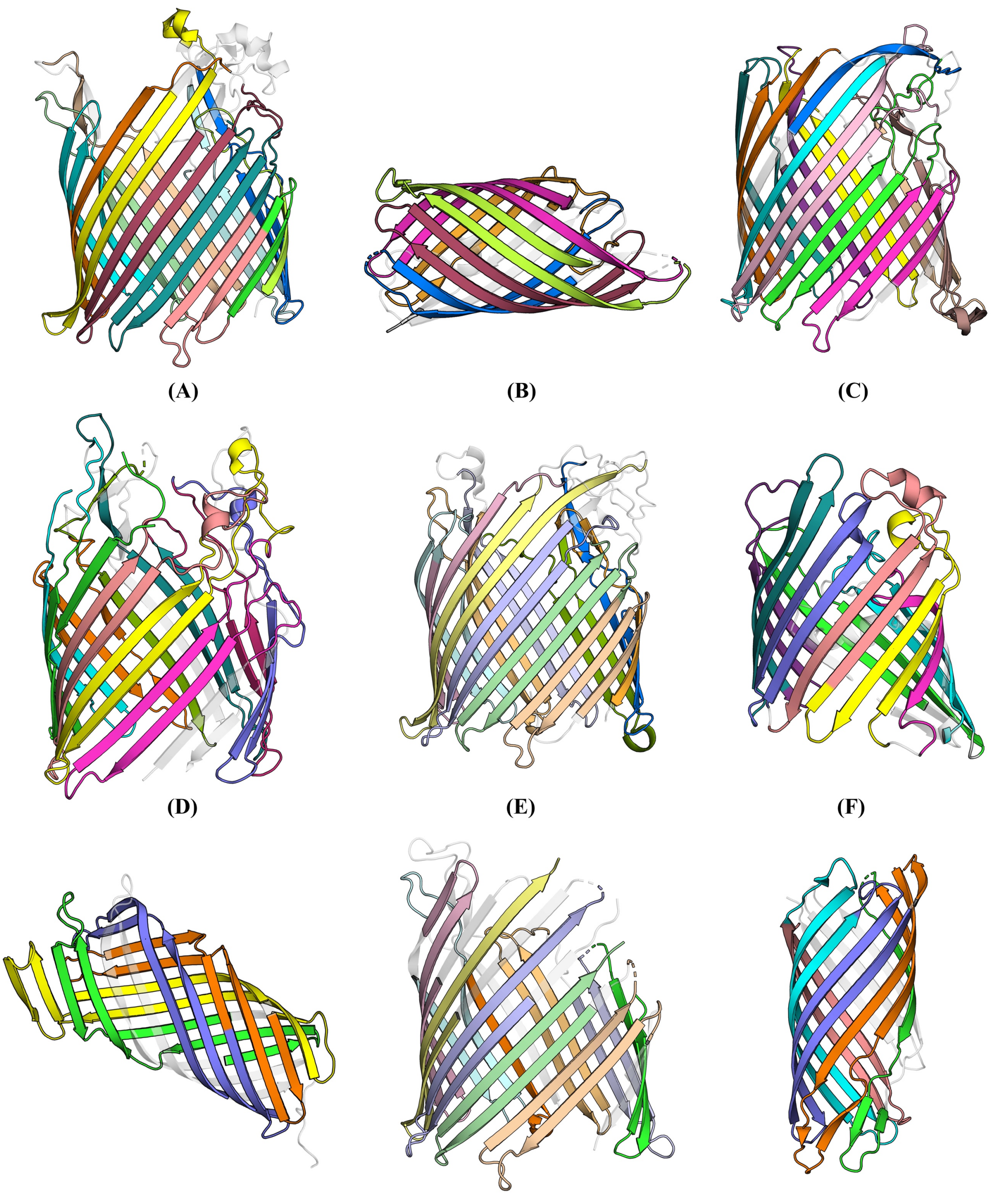

(A) (B) (C) (D) (E) (F) (G) (H) (I)

**Supplementary Figure 3:** A selection of β-barrel domains correctly detected by the pLM-Repeat method. Protein structures are colored according to repeat ranges. (A) 6E4V_A, (B) 2QOM_A, (C) 3QLB_A, (D) 1FEP_A, (E) 3ODW_A, (F) 6ENE_A, (G) 2X4M_A, (H) 3EFM_A, and (I) 4MEE_A.

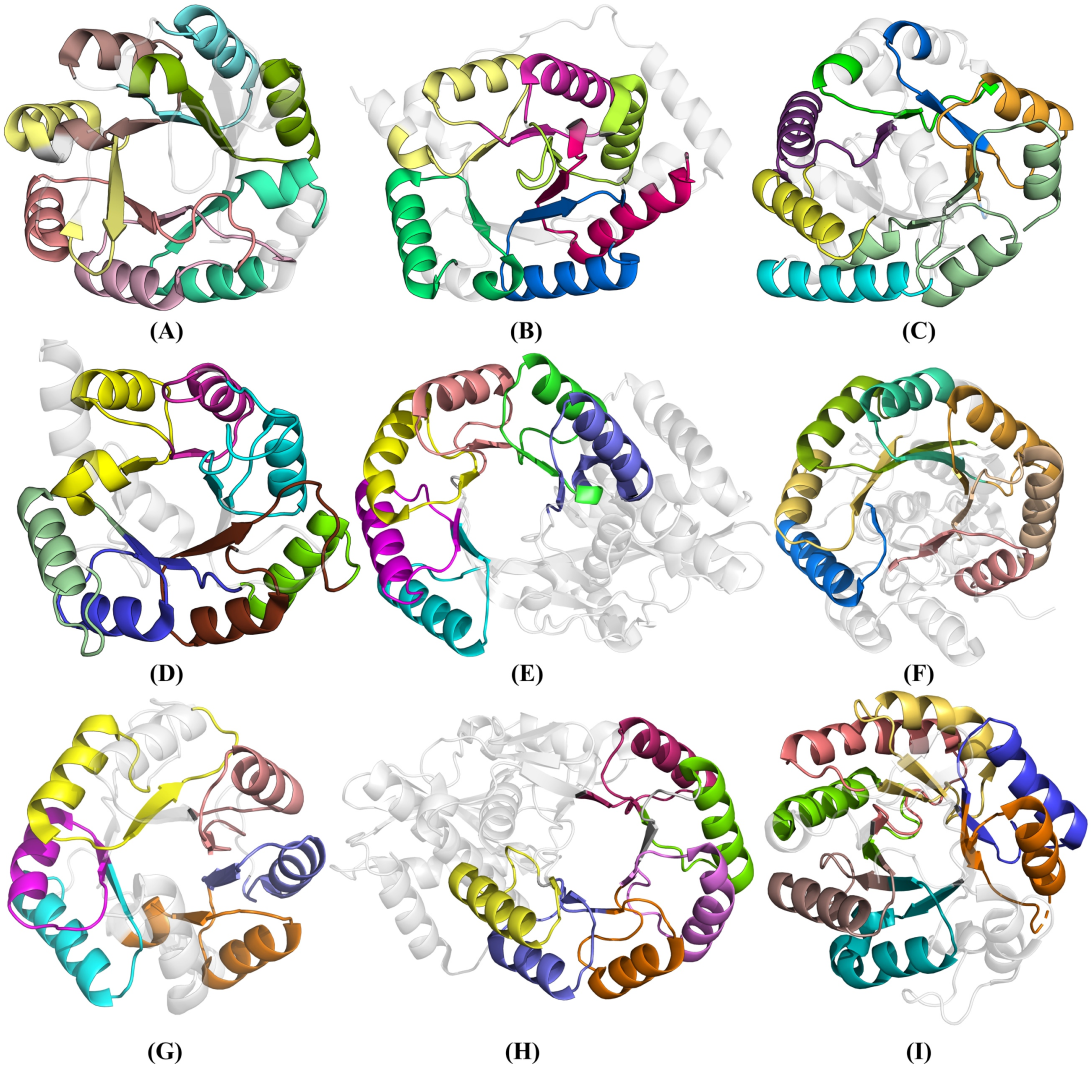

(A) (B) (C) (D) (E) (F) (G) (H) (I)

**Supplementary Figure 4:** A selection of TIM-barrel domains correctly detected by the pLM-Repeat method when the self-alignment score threshold was lowered to 0.25. Protein structures are colored according to repeat ranges. (A) 4W9T_A, (B) 3OA3_A, (C) 1A50_A, (D) 1YXY_A, (E) 2PGW_A, (F) 1NVM_A, (G) 3IGS_A, (H) 2QGY_A, and (I) 3JUG_A.

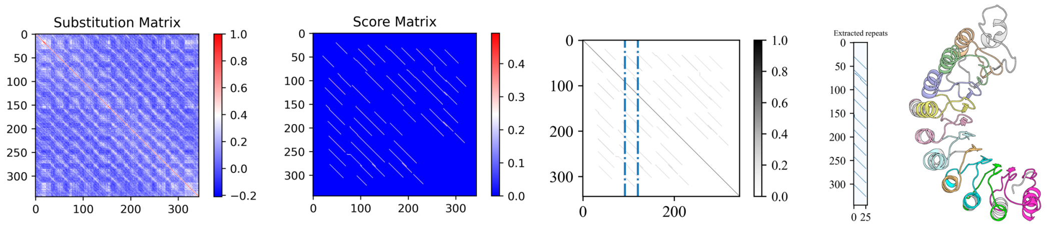


**Supplementary Figure 5:** pLM-Repeat intermediate outputs and detected repeats with ESM-IF structural embeddings as inputs on the domain 1K5D_C.

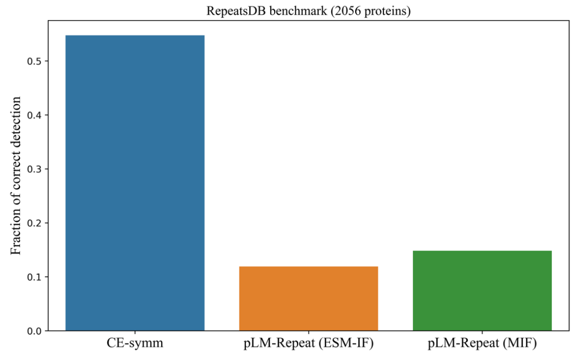


**(A)**

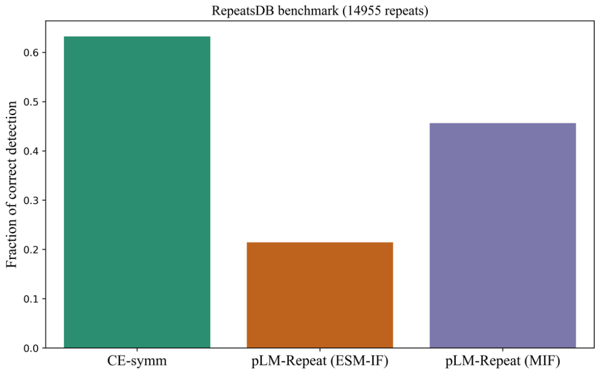


**(B)**

**Supplementary Figure 6:** Performance of pLM-Repeat equipped with structure embeddings (from two inverse folding models, ESM-IF and MIF) at the protein (A) and repeat (B) level, together with the state-of-the-art structure-based repeat detection software CE-symm.

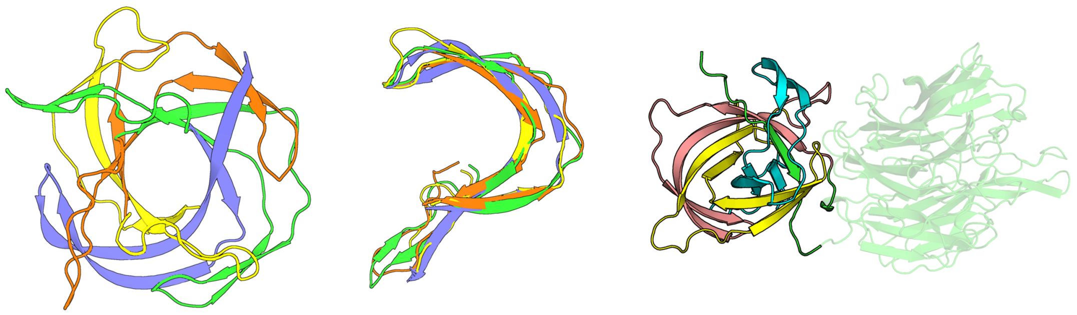

**(A)** **(B)**

**Supplementary Figure 7:** (A) The structure of domain A0A7C3HQW7 colored by four repeats. (B) A homolog (UniProt ID: A0A534S4A8) with three repeats.

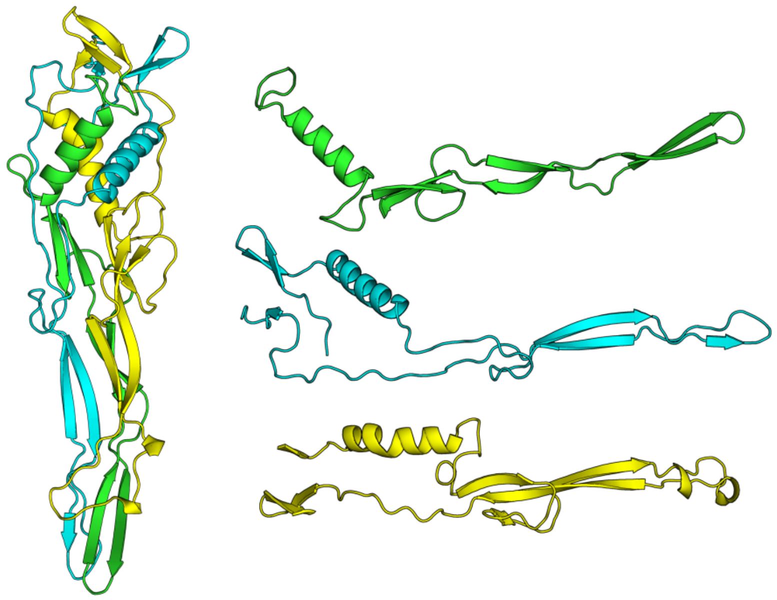

**Supplementary Figure 8:** The structure of domain A0A424SVE7 colored by three repeats.

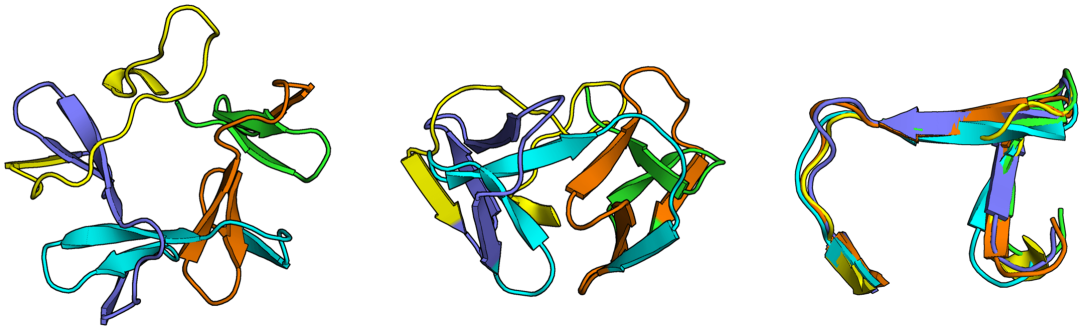

**Supplementary Figure 9:** The structure of domain A0A0S8GK70 colored by five repeats.

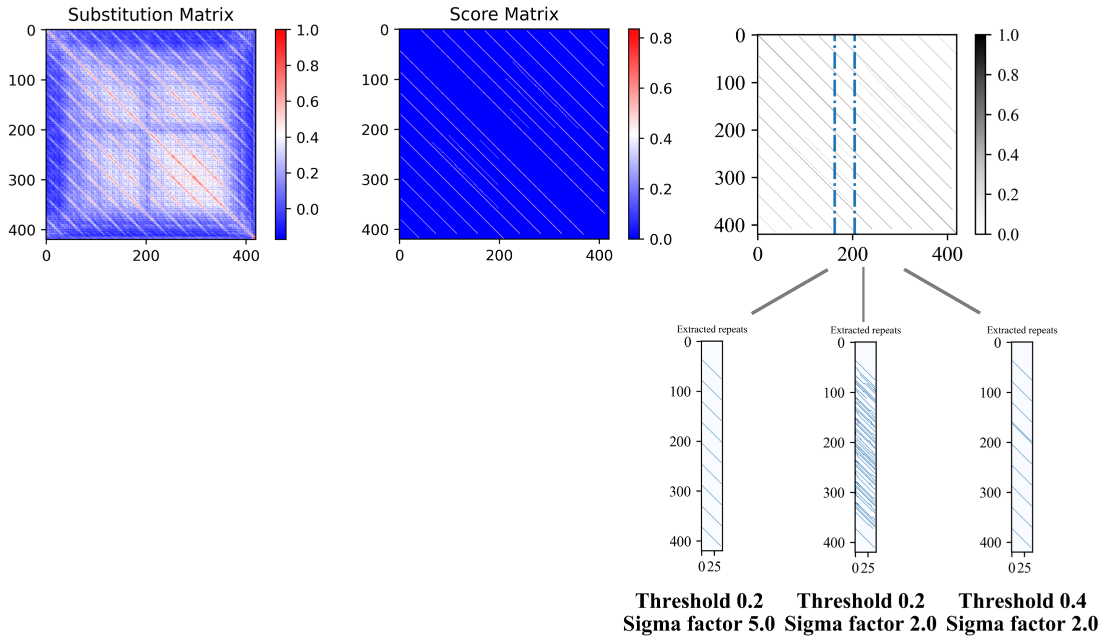


**Supplementary Figure 10:** The repeat extraction step is influenced by several factors, as in the example case of the domain PDB:4RV1_A. Substitution matrix, score matrix and sliding window are generated correctly. However, a Sigma factor of 2.0 and an extraction score threshold of 0.2 result in redundant detections. Increasing the sigma factor or the extraction score threshold helps to solve this problem and allows for clean detection of repeats.

**Supplementary Table 1:**

| Fold Class | pLM-Repeat (0.3) | pLM-Repeat (0.25) |
|---|---|---|
| β-Solenoid | 71 | 67 |
| α/β-Solenoid | 104 | 103 |
| α-Solenoid | 336 | **344** |
| β-Hairpins | 17 | 16 |
| Box | 40 | **43** |
| TIM-Barrel | 45 | **107** |
| β-barrel/hairpins | 41 | **45** |
| β-Propeller | 259 | **274** |
| α/β-Prism | 22 | **25** |
| α-Barrel | 24 | 23 |
| α/β-Trefoil | 38 | 36 |
| Aligned-prism | 7 | 6 |
| α-Beads | 7 | 7 |
| β-Beads | 18 | 16 |
| α/β-Beads | 18 | 18 |
| β-Sandwich-beads | 16 | 15 |
| α/β-Sandwich | 12 | **14** |

Number of correctly detected proteins in the RepeatsDB dataset based on fold classes in pLM-Repeat with a self-alignment score threshold of 0.25 and 0.3.
